# Development of a dedicated Golden Gate Assembly platform (RtGGA) for *Rhodotorula toruloides*

**DOI:** 10.1101/2022.03.02.482697

**Authors:** Nemailla Bonturi, Marina Julio Pinheiro, Paola Monteiro de Oliveira, Eka Rusadze, Tobias Eichinger, Gintare Liudžiūtė, Juliano Sabedotti De Biaggi, Age Brauer, Maido Remm, Everson Alves Miranda, Rodrigo Ledesma-Amaro, Petri-Jaan Lahtvee

## Abstract

*Rhodotorula toruloides* is a potential chassis to microbial cell factories as this yeast can metabolise different substrates into a diverse range of natural products, but the lack of efficient synthetic biology tools hinders its applicability. In this study, the modular, versatile and efficient Golden Gate DNA assembly system was adapted to the first basidiomycete, an oleaginous yeast *R. toruloides* (RtGGA). *R. toruloides* CCT 0783 was sequenced, and used for the RtGGA design. The DNA fragments were assembled with predesigned 4-nt overhangs and a library of standardized parts was created containing promoters, genes, terminators, insertional regions, and resistance genes. The library was combined to create cassettes for the characterization of promoters strength and to overexpress the carotenoid production pathway. A variety of reagents, plasmids, and strategies were used and the RtGGA proved to be robust. The RtGGA was used to build three versions of the carotenoid overexpression cassette were built by using different promoter combinations. The cassettes were transformed into *R. toruloides* and the three new strains were characterized. Total carotenoid concentration increased by 41%. The dedicated GGA platform fills a gap in the advanced genome engineering toolkit for *R. toruloides*, enabling the efficient design of complex metabolic pathways.

## 1 Introduction

The Sustainable Development Goals (SDGs) of the United Nations (UN) mark its historic shift towards the integration of economic and social development with environmental sustainability. Promoting food security and improved nutrition, ensuring sustainable use of resources and production patterns, and taking urgent action to combat climate change are among its directives(United Nations General Assembly, 2015). Microbial cell factories are an essential element for meeting UN’s SDGs as biofuels, pharmaceuticals, and other chemicals can be produced from sustainable raw materials using engineered or native organisms. Microbial cell factories do not compete with food and arable land resources, they are not directly affected by different locations or climates (Koutinas et al., 2014) and they can use lignocellulosic biomass hydrolysates and biological waste as substrates. Being able to use such substrates is interesting, due to their abundance, low cost, and sustainability (Adrio, 2017).

Strain development is a crucial part of improving titers, yields, and rates when using microbial cell factories and this work is typically onerous (involves millions of dollars), takes 3–10 years to complete (Opgenorth et al., 2019), and requires several loops of the Design-Build-Test-Learn (DBTL) cycle (Nielsen and Keasling, 2016). Developing versatile, fast, standardized, and modular tools for building pathways are necessary for obtaining optimized strains using the DBTL cycle (Larroude et al., 2019). There has been an advancement in synthetic biology in the past decade that has led to the creation of several robust gene assembly platforms that can be exploited for metabolic engineering and pathway optimization, such as Sequence and Ligation-Independent Cloning (SLIC) (Li and Elledge, 2007), Gibson assembly (GA) (Gibson et al., 2009), Circular Polymerase Extension Cloning (CPEC) (Quan and Tian, 2011), and Golden Gate assembly (GGA) (Engler et al., 2009, 2008).

GGA provides a way to construct, diversify, and optimize multi-gene pathways exploiting the ability of Type IIS restriction endonucleases, such as BsaI, to cleave outside of their recognition sequence. The recognition sites are strategically placed distal to the cleavage site of inserts and cloning vectors, and, as such, the endonucleases can remove the recognition site from the assembly leaving 4 nucleotides long overhangs. As the overhang sequence is not dictated by the endonuclease, no scar sequence is introduced, and the specificity of each overhang allows simultaneous orderly assembly of multiple fragments. Digestion and ligation can be carried out at the same time when combining a DNA ligase with the endonucleases (Engler et al., 2009, 2008), therefore, several DNA fragments can be seamlessly assembled in the desired order in a one-pot reaction (Weber et al., 2011).

The Yeast Toolkit (YTK) is based on GGA and relies on an organizational system of up to three tiers of plasmids for storage or use (Lee et al., 2015). Parts, such as promoters, genes, terminators, etc., are stored in level 1 plasmids, called parts library. At level 2, level 1 promoter, gene, and terminators are assembled to create transcriptional units (TUs). Multiple TUs and other parts (integration sites and markers) are assembled at level 3 into a multi-gene vector. The standardization of DNA fragments by position-specific 4 nt overhangs offers the opportunity to shuffle around parts in a one-pot reaction, making the diversification and optimization of engineered pathways a less laborious affair. YTK was developed for the most traditional yeast cell factory, *Saccharomyces cerevisiae*, but it has been expanded to other microbial cell factories, such as *Yarrowia lipolytica* (Celińska et al., 2017; Larroude et al., 2019), *Pichia pastoris* (Prielhofer et al., 2017), *Kluyveromyces marxianus* (Rajkumar et al., 2019), and *Ashbya gossypii* (Ledesma-Amaro et al., 2018).

Among the microbial cell factories, the nonconventional yeast *Rhodotorula toruloides* is considered a promising biotech workhorse (Park et al., 2017), as it can consume a variety of carbon and nitrogen sources (Lopes et al., 2020a; Xu and Liu, 2017), has an improved tolerance towards inhibitors present in lignocellulosic hydrolyzates (Bonturi et al., 2017a), and can naturally co-produce industrially important molecules (lipids, carotenoids, and enzymes) (Park et al., 2017). Microbial lipids can be considered potential feedstock for oleochemicals and applied in different products, such as biofuels, cosmetics, plastics, coatings, surfactants, lubricants, paints, among others (Adrio, 2017). Carotenoids are used for food and pharmaceutical industries as A-vitamin precursors, colourants, or antioxidants. *R. toruloides* produces mainly γ-carotene, β-carotene, torulene, and torularhodin (Mata-Gómez et al., 2018). Today, torulene and torularhodin are not as commercially important as β-carotene, but due to their valuable antioxidant properties, the interest in their production has been increasing lately (Kot et al., 2018).

In the past decade, omics studies shed light on the genome and metabolic pathways of *R. toruloides* (Hu and Ji, 2016; Kumar et al., 2012; Morin et al., 2014; Zhu et al., 2012), and its modification was made possible by the identification of constitutive (Liu et al., 2016; Wang et al., 2016) and inducible promoters (Johns et al., 2016; Liu et al., 2015), and selectable markers based in auxotrophies (Yang et al., 2008) and antibiotics (Lin et al., 2014). Despite these advances, there is still a lack of a high throughput DNA assembly platform for this yeast. This is attributed to several factors. *R. toruloides* genes have a very high GC content (∼62%) (Sambles et al., 2017a), which hinders PCR reactions due to disruption polymerase efficiency and accuracy, hinders gene synthesis, and the design specific primers without reaching an extremely high annealing temperature, and heterologous genes need to be codon-optimised (Lin et al., 2014). Also, there is a lack of known autonomously replicating sequence (ARS) elements, therefore metabolic engineering requires the integration of genes and pathways into the genome (Tsai et al., 2017). Furthermore, integration cassettes require long homology arms (500-1000 bp), as the non-homologous end joining (NHEJ) is the default mechanism of DNA repair in *R. toruloides*. Due to the latter, deleting the gene KU70, part of the main component of NHEJ, improves gene deletion and integration frequency without negatively affecting the oleaginous and fast-growing features of *R. toruloides* (Koh et al., 2014).

A crucial part of utilizing *R. toruloides* as a microbial cell factory is the ability to engineer the metabolic pathways in the cell in an efficient and standardized manner (Park et al., 2017). The current work aimed at the development of a YTK-dedicated *R. toruloides* Golden Gate assembly (RtGGA) platform. For this purpose, we have: (i) sequenced the genome of *R. toruloides* CCT0783, which derived the hydrolysate-tolerant strain CCT7815(Bonturi et al., 2017a; Lopes et al., 2020b; Pinheiro et al., 2020); (ii) characterized six native promoters; and (iii) as the proof of concept of such a platform, simultaneous deletion of KU70 and overexpression of three native genes in the carotenoid biosynthesis pathway – geranylgeranyl diphosphate synthase (crtE, EC 2.5.1.29), phytoene dehydrogenase (crtI, EC 1.3.99.30), and phytoene synthase/lycopene cyclase (crtYB, EC 2.5.1.32/EC 5.5.1.19) – was carried out. Once this platform was established, it was used for the assembly of different constructs to the metabolic engineering of *R. toruloides* for further improvement of the carotenoid production using xylose, the main sugar of hemicellulosic biomass hydrolysates.

## 2 Materials and Methods

### 2.1 Microorganism, genomic DNA extraction, sequencing, analysis, and annotation

The strain *R. toruloides* CCT 0783 was purchased from Coleção de Culturas Tropicais (Fundação André Tosello, Campinas, Brazil; synonym IFO10076). The yeast was reactivated and stored as described previously (Lopes et al., 2020a). The genomic DNA was extracted according to Otero et al. (2010). The DNA sequencing libraries were prepared with Illumina TruSeq Nano DNA Library Prep kit according to the manufacturer’s recommendations. The libraries were sequenced on MiSeq v3 flow cell, read length 2 × 175 bp using the MiSeq platform (Illumina Inc., San Diego, CA, USA). The library preparation and sequencing were performed in the Institute of Genomics Core Facility, University of Tartu. Reads were cleaned and trimmed with Trimmomatic-0.39 (Bolger et al., 2014). Draft assembly was created with spades-3.12.0 (Bankevich et al., 2012) using diploid mode. Protein coding sequences were determined from received haplocontigs by homology search with blastn (Altschul, 1997) against reference NP11 gene sequences with coverage and identity thresholds 80% and 70%, respectively. This Whole Genome Shotgun project has been deposited at DDBJ/ENA/GenBank under the accession JABGON000000000. The version described in this paper is version JABGON010000000. For creating the native library parts and yeast transformations, the strain *R. toruloides* CCT7815 was used. This strain is derived from *R. toruloides* CCT0783 after a short-term adaptation in sugarcane bagasse (Bonturi et al., 2017a).

### 2.2 Plasmids

The backbone plasmid used for storing level 1 parts was either pSB1K3-RFP from the iGEM collection (http://parts.igem.org/Collections) or TOPO™ Vector (Thermo Fisher Scientific, Carlsbad, USA). For level 2 and level 3 plasmids, pSB1C3-RFP from the iGEM collection. When the GGA reactions using iGEM collection plasmids did not work, pGGA (New England Biolabs, New England Biolabs, Ipswich,) containing a red fluorescent protein (RFP, BBa_E1010, http://parts.igem.org/Part:bba_E1010) (Celinska et al., 2017) was used instead. pSB1K3-RFP and TOPO vector carry resistance to the kanamycin gene, while pGGA and pSB1C3-RFP carry chloramphenicol resistance gene.

### 2.3 DNA sequences

The primer sequences used for removing internal BsaI cutting sites as well as to amplify all parts with their respective overhangs for the GGA can be found in Supplementary Table 1. The genomic DNA from *R. toruloides* was extracted (Lõoke et al., 2011) and used as a template for the amplification of insertional regions, promoters, terminators, and genes. For the insertional region, five hundred base pairs fragment upstream (insUP) and downstream (insD) the KU70 (DNA-dependent ATP-dependent helicase subunit 70, RHTO_06014) were amplified from the genome. The promoters from the genes glyceraldehyde-3-phosphate dehydrogenase (GAPDH, also widely named as GPD1 in *R. toruloides*’
ss literature, RHTO_03746), alcohol dehydrogenase 2 (ADH2, RHTO_03062), xylose reductase (XYL1, RHTO_03963), glucose 6-phosphate isomerase (PGI, RHTO_04058), and 1,6-bisphosphatealdolase (FBA, RHTO_03043) (Díaz et al., 2018; Wang et al., 2016). The intronic promoter from the gene LDP1 (lipid droplet protein 1 gene) (Liu et al., 2016) was synthesized (Twist Biosciences, San Francisco, USA) without BsaI recognition sites. The genes geranylgeranyl diphosphate synthase (crtE, RHTO_02504), phytoene dehydrogenase (crtI, RHTO_04602), and phytoene synthase/lycopene cyclase (crtYB, RHTO_04605) were amplified and used for proof of concept aiming at improving carotenoids production. BsaI recognition sites from the aforementioned genes were removed by inserting point mutations without changing the aminoacids by overlapping PCR. For terminators, the sequences from cauliflower mosaic virus 35S (t35S, GenBank No. MF116009) and the *Agrobacterium tumefaciens* nopaline synthase (tNOS, MF116010) (Fillet et al., 2017) were synthesized (Twist Biosciences). Native terminators from heat shock 70 kDa protein 1/8 (tHSP, RHTO_07842) and the GPD1 (tGPD) were amplified from the genomic DNA. The geneticin (G418), hygromycin (HYG), nourseothricin (NAT), and bleomycin (BLE) resistance genes were codon-optimised for *R. toruloides* using Benchling and synthesized either by IDT (Leuven, Belgium) or Twist Biosciences. When GC content was too high for synthesis and it was not possible to lower it without drastically changing the codons, the sequences were split into fragments for synthesis and assembled by either GGA or by overlapping PCR. The resistance genes were further assembled by GGA with XYL1 promoter (pXYL1) and tNOS, herein called marker (M).

### 2.4 DNA amplification by polymerase chain reaction (PCR)

DNA sequences below 3 kb were amplified via PCR using DreamTaq Green PCR Master Mix (2X) (Thermo Fisher Scientific, Vilnius, Lithuania) or HOT FIREPol GC Master Mix (Solis BioDyne, Tartu, Estonia), otherwise, the high-fidelities Phusion DNA Polymerase or Platinum SuperFi Master Mix (Thermo Fisher Scientific, Vilnius, Lithuania) were used instead. All reactions were set up and performed according to the manufacturer’s instructions for a high GC content template. Following PCR, the samples were loaded on a 1% agarose gel and the electrophoresis at 120 V. The fragments sizes were estimated using gene ruler 1 kb Plus DNA Ladder (Thermo Fisher Scientific). Amplicons with the correct size were excised from the gel and purified using Favorprep™ GEL/PCR Purification Kit (Favorgen, Wien, Austria) following the manufacturer’s instructions, quantified by Nanodrop (Thermo Fisher, Whaltan, USA), and used for GGA assembly and TOPO subcloning.

### 2.5 Construction of pGGA carrying a red fluorescence protein (RFP)

To facilitate the screening of bacterial colonies containing GGA constructs, a variant of pGGA containing the RFP flanked by overhangs and BsaI cutting sites was constructed. Both pGGA and RFP were amplified by PCR, purified, and assembled by Gibson Assembly (Gibson et al., 2009) reaction according to the manufacturer’s instructions (Gibson Assembly Master Mix, New England Biolabs, Ipswich, USA). Two microliters of the assembly reaction were taken for the consequent bacterial transformation.

### 2.6 Assembly of levels 1, 2, and 3 constructs by GGA

Level 1 (Lv1, Figure 1A) comprised the storage of single standardized parts (promoters, genes, markers, insertional regions, and terminators) in a plasmid. Assembly reactions were set up in the following manner: 75 ng of pSB1K3 and 150 ng of insert purified after electrophoresis, 2 µl of T4 DNA Ligase Buffer (New England Biolabs), 1 µl of T4 or T7 DNA Ligase (New England Biolabs), 1 µl of BsaI-HFV2 (New England Biolabs), up to 20 µl of nuclease-free H_2_O. The mixture was incubated at 37°C for 1 h, followed by 5 min at 60°C. This approach was used for establishing the initial library (RtGGA v1.0, Figure 1A) needed for constructing the pathway overexpressing the native carotenoids pathway as a proof of concept. The expansion of the GGA library used the TOPO™ XL-2 Complete PCR Cloning Kit (Thermo Fisher Scientific, Vilnius, Lithuania) instead as it allowed the storage of the part maintaining the flanking BsaI cutting sites (RtGGA v1.1, Figure 1A).

**Figure 1.**
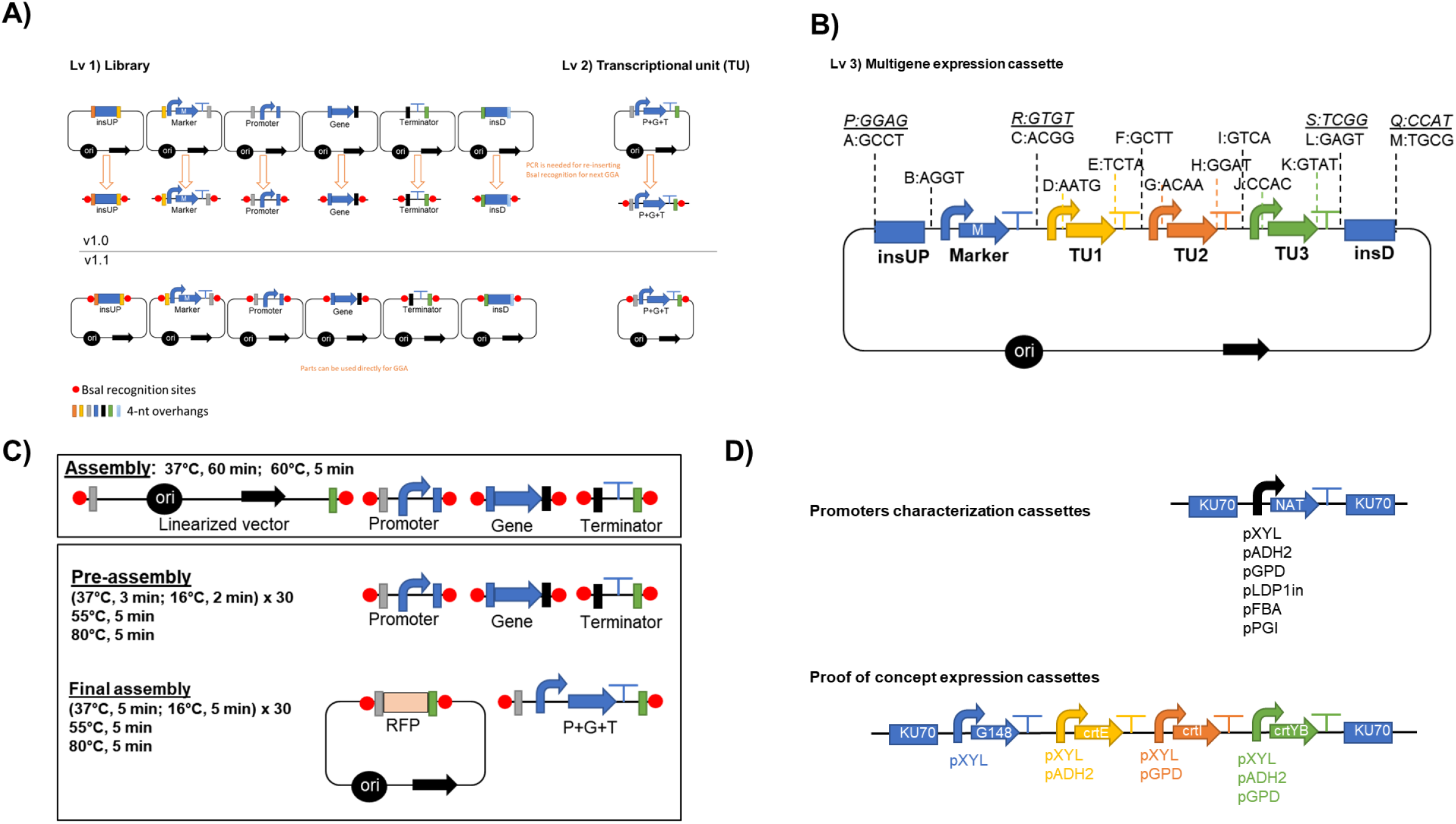
The RtGGA with **A)** Level 1 (Lv1) for the library of standardized parts−containing regions upstream (insUP) and downstream (insD) an insertional site, selection marker (M), promoters, genes, and terminators− and level 2 (Lv 2) by combining promoter (P), gene (G), terminator (T) into transcriptional units (TU). The upper part illustrates version 1.0 of Lv1 and Lv2, while the lower part illustrates version 1.1 of the RtGGA which BsaI recognition sites are flanking the parts and TUs to enable direct use of them into GGA reactions. **B)** Level 3 (Lv3) RtGGA multigene expression cassette and standardized overhangs from the *Y. lipolytica* GGA platform (Celińska et al., 2017). The overhangs in *italic* are the changes made to make it compatible with *R. toruloides* **C)** Upper box shows the reaction method proposed in the *Y. lipolytica* GGA platform (Celińska et al., 2017) for level 2 assembly using linearized PCR-amplified parts. The lower box shows the two-step assembly protocol adapted from Larroude et al. (2019) for the assembly of Lv2. **D)** Scheme of the cassettes assembled for the characterization of the promoters strength and the overexpression of the carotenoid pathway. Legend: ori – the origin of replication; black arrow– bacterial resistance gene; RFP: red fluorescence protein;

The level 2 (Lv2, Figure 1A) of GGA was the assembly of a gene (G) with a promoter (P) and terminator (T), resulting in a transcriptional unit (TU) (Figure 1B). For the proof of concept of the GGA platform, the genes crtE (G1), crtI (G2), and crtYB (G3) were combined with three different promoters (pXYL1, pGPD1, and pADH2), and tNOS using the pSB1C3 or pGGA plasmid as vectors. A total of nine Tus were generated. The reactions were done using 150 ng of each part of the TU and either used the same recipe as level 1 or by using the NEB Golden Gate Assembly Mix (New England Biolabs) according to the manufacturer’s instruction. A two-step assembly protocol adapted from Larroude et al. (2019) was also tested (Figure 1B).

The final step of the GGA platform is to assemble the pathway with a selection marker and the insertional regions for guiding the pathway integration with simultaneous deletion of a target gene (Figure 1C). This structure was used for assembling both the cassette for the characterization of the strength of promoters from the library (pXYL1, pGPD1, pFBA, pADH2, pLDP1, and pPGI) and the overexpression of the carotenoid pathway (RtGGA proof of concept). The promoter strength was evaluated in terms of the expression of NAT gene as done similarly by Wang et al. (2016). Lv1 parts of KU70 insUP and insD, NAT, tNOS, and promoters were assembled (Figure 1D) by RtGGA in pGGA by reaction conditions described for library assembly.

For the proof of concept, G418 marker was combined with the 3 TUs flanked by the insertional region aiming at the deletion of KU70 gene (Figure 1B and D). Three different level 3 constructs were obtained by using different combinations of promoters (Table 1). This was done by PCR amplification of Lv1 and Lv2 the aforementioned parts followed by subcloning them into PCR-XL-2-TOPO™ Vector (Thermo Fisher Scientific). The reaction for TOPO™ Cloning was modified as follows: 1 µl of PCR-XL-2-TOPO™ Vector, insert required for 5:1 molar ratio of insert:vector, 1 µl of Salt Solution, up to 6 µl of deionized H_2_O. The mixture was incubated at 25°C for 1 h. Two microliters of the cloning reaction were used for bacterial transformation. The reaction to assemble level 3 (Lv 3) GGA was prepared as follows: 75 ng of pGGA, the mass of each insert (TOPO™ plasmid) required for 2:1 molar ratio of insert:vector, 2 µl of T4 DNA Ligase Buffer (New England Biolabs), 1 µl of T4 or T7 DNA Ligase (New England Biolabs, Ipswich, USA), 1 µl of BsaI-HFV2 (New England Biolabs), up to 20 µl of nuclease-free H_2_O. The reaction was performed as instructed by the official NEB GGA kit (37°C for 5 min followed by 16°C for 5 min) x 30 and 60°C for 5 min.

**Table 1.** Yeast strains created by the integration of the different constructs done with RtGGA in *R. toruloides* CCT7815.

| Strain number | Description |
| --- | --- |
| SBY92 | $\Delta ku70:: pXYL-G418-tNOS-pXYL-crtE-tNOS-pGPD-crtI-tNOS-pADH2-crtYB-tNOS$ |
| SBY93 | $\Delta ku70:: pXYL-G418-tNOS-pADH2-crtE-tNOS-pGPD-crtI-tNOS-pXYL-crtYB-tNOS$ |
| SBY94 | $\Delta ku70:: pXYL-G418-tNOS-pADH2-crtE-tNOS-pXYL-crtI-tNOS-pGPD-crtYB-tNOS$ |
| SBY109 | $\Delta ku70:: pADH2-NAT-tNOS$ |
| SBY116 | $\Delta ku70:: pLDP1in-NAT-tNOS$ |
| SBY117 | $\Delta ku70:: pXYL-NAT-tNOS$ |
| SBY119 | $\Delta ku70:: pGPD-NAT-tNOS$ |
| SBY124 | $\Delta ku70:: pPGI-NAT-tNOS$ |

All assemblies from GGA and TOPO™ subcloning were used for bacterial transformation using *Escherichia coli* DH5α according to the manufacturer’s instructions. White colonies were screened by PCR using SP6/T7 and M13F/M13R primers (Supplementary Table 1), respectively. Colonies bearing constructs with the correct size were grown overnight using 4 ml of LB medium (10 g/l tryptone, 5 g/l yeast extract, 10 g/l NaCl) and the selection antibiotics. Plasmids containing the assemblies were recovered using Favorprep™ Plasmid DNA Extraction Mini Kit (Favorgen, Wien, Austria). DNA Sanger sequencing was used to confirm parts and constructs.

### 2.7 Yeast transformation and screening of correct transformants

*R. toruloides* was transformed according to the protocol described by Nora et al., (2019) with either the different level 3 constructs for the proof of concept or promoter strength characterization. *R. toruloides* CCT7815 (parental strain) was inoculated in 10 ml of YPD medium (glucose, 20 g/l; yeast extract, 10 g/l; peptone, 20 g/l) and incubated overnight with a stirring speed of 160 rpm at 30 °C. The culture was diluted to an OD600 of ∼0.2 in 10 ml of YPD and incubated again until an OD600 of ∼0.8. The culture was harvested in a sterile 50 ml centrifuge tube at 3,000 *g* for 10 min. The cell pellet was resuspended in 25 ml of sterile water and centrifuged again. The cells were resuspended in one milliliter 100 mM lithium acetate (LiAc) (pH 7.5) and 500 µl of the suspension was transferred to a 1.5 ml microfuge tube. The cells were centrifuged at 3,000 *g* for 30 seconds and the LiAc was removed. In the following order, 240 µl of polyethylene glycol (PEG) 4000 (50% w/v), 36 µl of 1M LiAc (pH 7.5), 24 µl of sterile water were added. Ten µl of salmon sperm DNA (10 mg/ml, pre-boiled at 100 °C for 10 min) was transferred to the tube before adding 50 µl of transforming DNA (0.1-10 μg, removed from the bacterial backbone by either restriction enzymes or by PCR amplification). The cell pellet and the added reagents were mixed completely by vigorous vortexing. After incubating the tube at 30 °C for 30 min, 34 µl of dimethyl sulfoxide (DMSO) was added to the mixture, which was then heat-shocked at 42°C for 15 min. The tube was centrifuged at 3000 *g* for 30 s and the supernatant was removed. The pellet was resuspended in 2 ml of YPD and transferred to a new 15 ml tube, which was incubated overnight with shaking at 30 °C. The following day the suspension was plated onto solid YPD containing selection pressure and left to grow at 30 °C for two days. For the strains transformed with level 3 carotenoids multigene expression cassette G418 was used, while for the promoter characterization NAT was used (100 mg/ml for pPGI and pXYL1, 50 mg/ml pADH2 and pFBA, 25 mg/ml pLDP1 and pGPD1).

For the screening of correct transformants, yeast colonies were checked by PCR and sequencing. The yeast colony was grown overnight in 10 ml of YPD and the gDNA was extracted using the Wizard Genomic DNA Isolation kit (Promega, Switzerland). The PCR reaction was as follows: 40 ng of template, 5 µl Platinum SuperFi II Master mix (Thermo Fisher Scientific, Vilnius, Lithuania), 2 µl of SuperFi GC enhancer (Thermo Fisher Scientific, Vilnius, Lithuania), 1 mmol/l each of reverse and forward verification primers (Supplementary Table S1), and water to 10 µl. Reaction conditions were done according to the manufacturer’
ss instructions. The amplicons were evaluated by electrophoresis and the ones with the correct size were purified from the agarose gel and sequenced.

### 2.8 Characterization of the strains in shake flasks

Xylose was used as a carbon source for the characterization of strains presented in Table 1. Mineral medium, containing, per liter, 30.0 g carbon source, 0.8 g (NH_4_)_2_SO_4_, 3.0 g KH_2_PO_4_, 0.5 g MgSO_4_·7 H_2_O, 1.0 ml vitamins solution, and 1.0 ml trace metal solutions were used (Lahtvee et al., 2017). The carbon to nitrogen (C/N) ratio of the medium was 80 (mol/mol). The cells were incubated at 30 °C and at a stirring speed of 160 rpm until xylose depletion. For all the strains, samples were taken for the yeast characterization in terms of growth (absorbance at 600 nm; OD600) and RNA extraction (mid-exponential phase). Strains *R. toruloides* CCT7815 and SBY92-94 were also characterized in terms of metabolites profile (HPLC) and production of carotenoids.

### 2.9 RNA extraction and cDNA synthesis

Materials used were RNase free and all steps were done at room temperature unless stated otherwise. One milliliter aliquots of the cell liquid culture from the mid-exponential phase (OD600 between 14-18) was collected and harvested at 23,000 *g* for 30 s at 4 °C. After centrifugation supernatant was quickly discarded and the cell pellet was frozen in liquid nitrogen. Samples were stored at -80 °C. RNA extraction was done using the Tri reagent kit (Invitrogen, USA). A few modifications were done to the protocol that was provided by the manufacturer. Before cell lysis in 1 ml of Tri reagent, the cell pellet was washed twice in 1 ml of RNAse free water using centrifugation as described above. One milliliter of Tri reagent was added to the biomass, which was resuspended by pipetting and incubated for 5 min followed by the addition of 200 µl of chloroform. The suspension was thoroughly mixed using a vortex, followed by 15 min of incubation. A triphasic system was obtained after centrifugation at 12,000 *g* for 15 min at 4 °C. The upper aqueous phase containing the RNA was transferred to a new tube and 0.5 ml of ice-cold 2-propanol was added. The mixture was incubated for 10 min and the RNA pellet was recovered after centrifugation at 12,000 *g* for 15 min at 4 °C and washed with 1 mL of ice-cold 75% (v/v) ethanol. After removing the supernatant, the RNA pellet was dried and 50 µL of RNA-free water was added for resuspension. RNA was kept at -80 °C until the cDNA synthesis which was done using the First Strand cDNA Synthesis Kit (Thermo Scientific, USA) according to the manufacturer’s instructions for GC-rich templates. Five hundred nanograms of RNA were used for the cDNA synthesis.

### 2.10 Measurement of relative expression of promoters and carotenogenic genes by real-time PCR

Relative gene expression by rtPCR was used to determine the strength of six native promoters and to validate the overexpression of the carotenogenic pathway. Promoter strength was measured by the expression of NAT gene and pADH2 was chosen as reference. For the assessment of the overexpression of native crtE, crtI, and crtYB, *R. toruloides* CCT7815 (parental strain) was chosen as a reference. GAPDH (glyceraldehyde 3-phosphate dehydrogenase) was used as an endogenous control (Bonturi et al., 2017). Three biological and five technical replicates were used. The resulting cDNA from section 2.9 was diluted 2.5 times right before use. The reaction was done as follows: 2 μl of HOT FIREPol® EvaGreen® qPCR Supermix (EG) (SolisBiodyne, Estonia), 3 mmol/l of each forward and reverse primer (Supplementary Table 1), 1 μl cDNA and Milli Q purified water to a final volume of 10 μl. Reaction conditions were: 95 °C for 12 min and 45 x (95°C for 15 s, 64°C for the 20 s, 72°C for 20 s). The gene relative gene expression was calculated as described in Bonturi et al. (2017).

### 2.11 Analytical methods

Microbial growth was estimated by measuring OD600 nm and converted to biomass concentration by using a calibration curve. Concentrations of metabolites, such as xylose, organic acids and glycerol were measured using high-pressure liquid chromatography (HPLC) (LC-2050C, Shimadzu, Kyoto, Japan) equipped with HPX-87H column (Bio-Rad, CA, USA) and a refractive index detector (RID-20A, Shimadzu, Kyoto, Japan), at 45 °C and 5 mmol/l H_2_SO_4_ as mobile phase with isocratic elution at 0.6 ml/min.

Once xylose was depleted completely the final cell biomass was recovered by centrifugation at 4,000 *g* for 10 min and washed twice with distilled water. Cells were frozen at -80 °C overnight and lyophilised. The final dry cell mass was determined gravimetrically.

Carotenoids were extracted from liophylised biomass as described in Pinheiro et al. (2020). Quantification was done by measuring absorbance at 450 nm (Larroude et al 2019) in a spectrophotometer and also by adapting to HPLC an ultra-performance liquid chromatography (UPLC) method described in Pinheiro et al. (2020). The separation was done at 40 °C in a C18 column (Kinetex 2.6 μm C18 100 Å, 100 × 4.6 mm, Phenomenex, Torrance, USA) using isocratic elution for 10 min using acetone:water (70:30, %v/v), followed by a gradient to 100% of acetone in 2 min, and re-equilibration for 5 min using acetone:water (70:30, % v/v). A flow rate of 0.8 ml/min was used. The UV detector was set to 40 °C and 450 nm. Detected peaks were identified according to the retention time profile determined by Weber (2007). β-carotene (Alfa Aesar, MA, USA) was used for the construction of the calibration curve for both methods.

## 3 Results and discussion

### 3.1 Genome sequence of *R. toruloides* CCT 0783

The genome of *R. toruloides* CCT0783 was sequenced in order to identify BsaI recognition sites and to allow the design of primers for the amplification of insertional regions, promoters, terminators and genes. llumina MiSeq platform was used, and the sequence was deposited (https://www.ncbi.nlm.nih.gov/genome/13280?genome_assembly_id=1549609). The total GC content was as expectedly high (61.9%) while the length of 40.59 Mb was determined. The determined length is about double the size of the genome of other *R. toruloides* strains deposited in NCBI (about 20 Mb; https://www.ncbi.nlm.nih.gov/genome/browse#!/eukaryotes/13280/). *R. toruloides* CGMCC 2.1609 was the only strain with an intermediate genome size of 33.39 Mb. About 55% of the genome of this strain posses ≥99% identity to the genome of haploid strain CBS 14 (A1 mating-type; synonym to IFO 0559) while 41.6% is highly similar to the strain IFO 0880 (A2 mating-type; synonym to CBS 349) (Sambles et al., 2017b). A similar finding was obtained with the genome of *R. toruloides* CCT 0783, where two versions of the same gene were found, one presenting ≥90% identity and the other version presenting ≥70% identity to the genome of haploid strain NP11. Further studies and analysis are being carried out to understand the ploidy, phylogenetic tree, and which of the versions of the genes are being expressed. Since the goal of this work was to develop a standardized, robust, and versatile platform for golden gate assembly for *R. toruloides* (RtGGA), the genomic DNA of the strain used in this work was used for designing primers, accessing the presence of BsaI recognition sites, and validating the constructs by Sanger sequencing.

### 3.2 Constructing standardized biological parts for RtGGA

The foundation of synthetic biology lies in standardized biological parts that would allow us to build new biological systems more efficiently and offer better designs for metabolic engineering approaches. The YTK for *S. cerevisiae*, a diverse collection of parts, exemplifies how crucial is standardization for synthetic biology. The YTK has a wide range of regulatory elements and selection markers, which makes engineering the genome and fine-tuning gene expression much less laborious. Expanding this technology to a wider range of microorganisms would allow swift metabolic engineering for wider biotechnology applications. Maintaining similar standards for assemblies is also helpful for exchanging parts between different yeasts.

With this goal in mind, we have developed a dedicated GGA platform for *R. toruloides*, based on the YTK standard and various parts collections developed for other yeasts (Celińska et al., 2017; Prielhofer et al., 2017; Rajkumar et al., 2019). Due to large differences in the genus of *R. toruloides* and other yeasts with dedicated GGA platforms, it was necessary to extensively optimize almost all steps of the assembly. *R. toruloides* has a GC-rich genome, leading to an outcrop of problems, from extremely high melting temperatures during primer design to the formation of secondary structures. One of the key changes that resulted in a successful assembly of the final construct was switching from linearized DNA parts (RtGGA v1.0) to plasmids (RtGGA v1.1), which allowed the restriction enzyme to anchor to the DNA more efficiently. Even while using the TOPO™ Cloning technology that had been developed for easy cloning, it was required to optimize the established method to accommodate the GC-rich genome of *R. toruloides*.

A dedicated GGA platform enables fast construction of complex expression cassettes for metabolic engineering in *R. toruloides*. A set of thirteen 4-nucleotide (nt) overhangs from the *Y. lipolytica* GGA system (Celińska et al., 2017) was initially used for this task. This set covered all three transcription units (TUs, containing P – promoter, G – gene, T - terminator), selection marker M (also a TU with P-G-T), and upstream and downstream insertional units, insUP and insD, respectively (Figure 1D).

For the assembly of level 1, the parts and the vector pSBIK3_RFP from the iGEM collection were used. For each position, the plasmid, along with the sequence to be used as the building block of the GGA, was PCR amplified using the corresponding primers containing the BsaI recognition site and the 4-nt overhang (Supplementary Table 1). In the case of a successful round of GGA reaction and transformation, the original red colonies of the pSBIK3_RFP plasmid were changed to white ones, as the GGA building block was inserted instead of the red fluorescence protein (RFP) reporter gene, therefore, allowing a quick and easy pre-screening.

The parts library (Figure 1A) was successfully assembled and contained: (i) six promoters (pGPD1, pXYL1, pADH2, pLDP1in, pPGI, pFBA); (ii) three native genes from the carotenogenic pathway (crtE, crtI, and crtYB); (iii) terminators (tNOS, t35S, tHS, tGPD); (iv) insertional regions, 500 bp upstream and downstream the KU70 gene, insUP and insD, respectively; and (v) resistance genes (G418, NAT, BLE, HYG). All parts were verified by DNA sequencing.

Level 2, TU, consisted of PCR amplifying parts from level 1 to re-insert the BsaI cutting sites and then to use the linear fragments for the GGA (Figure 1A, v1.0). In total, nine TUs were assembled by having all three carotenogenic genes under the three different promoters (Figure 1A). The TUs were assembled by using diverse approaches: (i) either using the official GGA kit or by using enzymes separately (BsaI and T4 or T7 ligase); (ii) different plasmids (pSB1C3_RFP or pGGA_RFP); and (iii) either by one or two-step protocol (Larroude et al., 2019).

The RtGGA showed to be a robust system regardless of the assembly strategy, backbone, and the usage of the official kit or not, but with the increasing assembly complexity and high-GC content DNA, a higher efficiency was obtained by using a two-step assembly protocol (Figure 1D). The two-step assembly method most likely avoids unspecific ligations between overhangs. All nine TUs were assembled effectively and confirmed by sequencing. Once the level 2 construction was successful, the cassette for promoter strength characterization (Figure 1D) was built and transformed in the parental strain (*R. toruloides* CCT7815). This assessment would further be used for selecting the promoters for assembling the multigene cassette.

### 3.3 Characterizing promoters for *R. toruloides*

As the *R. toruloides* CCT 0783 strain showed to have differences in terms of its genome size, and probably ploidy or in terms of expression of genes as two different versions of the same gene were identified, we decided to characterize the promoter strength in its derived strains. Six native promoters were selected based on previous works with *R. toruloides* (Liu et al., 2016; Wang et al., 2016; Díaz et al., 2018; Nora et al., 2019), assembled with NAT gene, tNOS and flanking regions (insUP and insD) targeting the KU70 loci for integration. The six different cassettes were transformed into *R toruloides* and the resulting strains were grown in minimal medium and xylose as carbon source. Biomass samples were collected from the exponential growth phase for the promoters strength characterization in terms of relative expression of NAT gene by using rtPCR. The promoter ADH2 was chosen as a control as its NAT expression showed the lowest standard deviation among the biological replicates. The order of strength in xylose-grown exponential cells was pGPD<pLDP1in<pADH2=pFBA<pXYL<pPGI (Figure 2). Díaz et al. (2018) also reported pXYL as being a stronger promoter than pGPD for *R. toruloides* CECT 13085 for xylose-cultivated cells regardless of the growth phase. For *R. toruloides* AS 2.1389 grew on YPD medium the promoter strength order was pGPD<pFBA<pPGI (Wang et al., 2016). One report that showed discrepancy is the one from Liu et al. (2016) that reported stronger activity of pLDP1in when compared to pGDP1 in glucose-grown *R. toruloides* ATCC 10657. Nora et al. (2019) characterized a wide range of native promoters for *R. toruloides* IFO0880 and pGPD1 was defined to be a medium-strong promoter. The authors also reported that pGDP1 showed less predictable expression levels than other promoters accessed, as the levels were different depending on the reporter gene and cultivation medium. Differences in promoter strength found in different literature might be explained by using different strains, media, and characterization methodologies, as some works measured expression by the fluorescence produced by expressing reporter proteins (Nora et al. 2019; Díaz et al. 2018; Liu et al., 2016) or by a reporter gene relative expression using RT-PCR (Wang et al., 2016). A weak (pGPD1), a medium (pADH2), and a strong (pXYL1) promoters were selected to be tested further in the proof of concept cassette.

**Figure 2.**
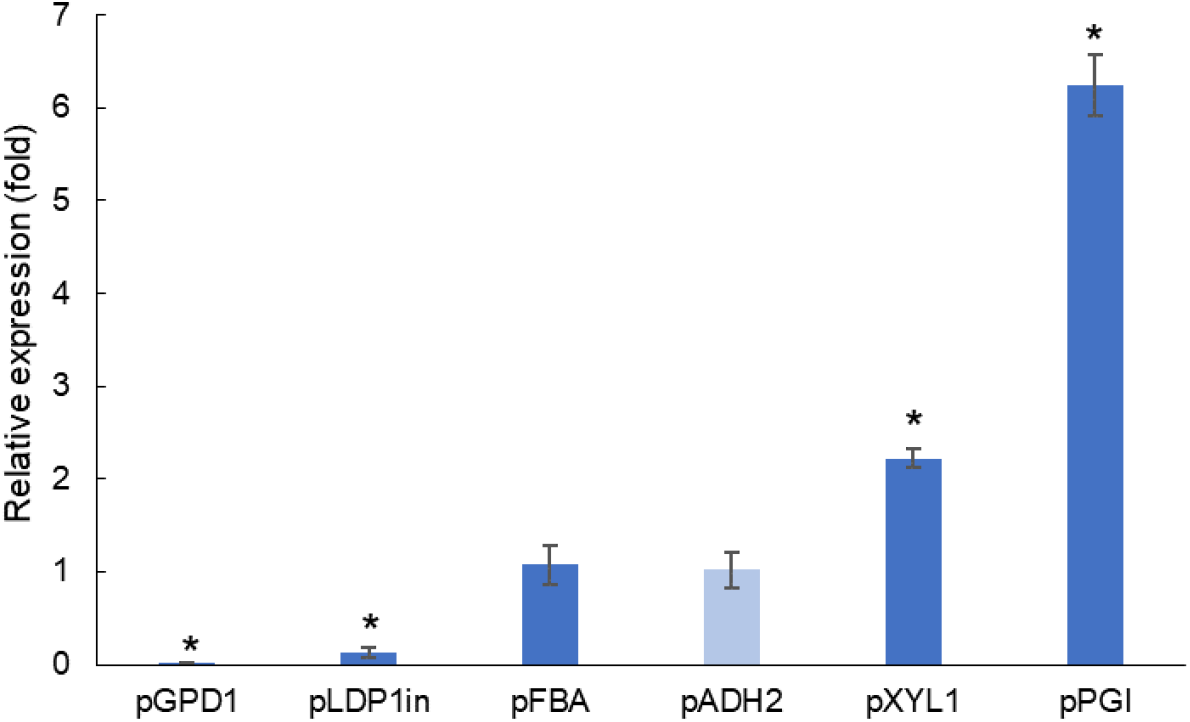
Relative expression in terms of the fold of the NAT gene under different promoters. pADH2 (light blue) was used as a reference and the asterisk indicates a difference with statistical significance (Student’
ss t-test; p-value ≤ 0.05).

### 3.4 Proof of concept of RtGGA for the overproduction of carotenoids

At first, for level 3 construction of the proof of concept cassette, it was required to assemble into GGA plasmid the following PCR amplified parts: insUP, Marker, TU1, TU2, TU3, and insD (RtGGA v1.0). The initial assembling of the level 3 construct turned out to be unsuccessful using GGA plasmid, while different concentrations of enzyme (BsaI and T7 or T4 DNA ligase) and buffer, different pre-assembly strategies, or even switching to the official NEB GGA kit were tested. To overcome the issue, new overhangs (C and L) were designed with no potential similarity with each other and, instead of using PCR amplified parts, all parts were subcloned to TOPO vectors with BsaI recognition sites and overhangs (Figure 1B). The subcloning of level 1 parts (insUP and insDO) worked by using the manufacturer’s protocol, but for subcloning the TUs and marker it was required to optimize the protocol. Different ratios of TU fragment to TOPO™ vector were used, and a 5:1 molar ratio was found to be the most efficient (data not shown). Three different level 3 constructs containing the crtE, crtI, and crtYB genes under different combinations of pXYL, pGPD, and pADH2, respectively, were assembled by GGA by the one-step reaction. Using parts inside plasmids probably increases the efficiency of BsaI recognition and cutting by offering a higher surface for the enzyme to dock on. With the increased success rate of using parts in plasmid form rather than linearized form, all parts from the library were also subcloned to TOPO vector bearing overhangs and BsaI recognition sites (RtGGA v1.1, Figure 1A). As recommendations, it is suggested to simulate *in silico* GGA to check for any possible warnings. For level 1 and level 2 constructs done from PCR-amplified fragments worked, but it is more convenient to blunt-end clone the amplicons and have ready-to-use parts for any type of level. In case of failure by using the manufacturer’
ss protocol it is suggested to use the two-step or other types of pre-assemblies. The efficiency of the developed GGA was validated by overexpressing the three most important genes in the carotenoids production pathway - crtE, which catalyzes the formation of an important precursor of carotenoids; crtI, and crtYB, both of which are involved in the production of γ-carotene, β-carotene, torularhodin and torulene (Figure 3). In this work, we aimed to validate the Rt-GGA by regulating the expression of these genes under different combinations of promoters, such as the strong xylose reductase (pXYL1), mild alcohol dehydrogenase 2 (pADH2), and weak glyceraldehyde 3-phosphate dehydrogenase (pGPD1). The constructions were transformed into *R. toruloides* CCT7815 and three new strains were obtained: SBY92, SBY93, and SBY94 (Table 1).

**Figure 3.**
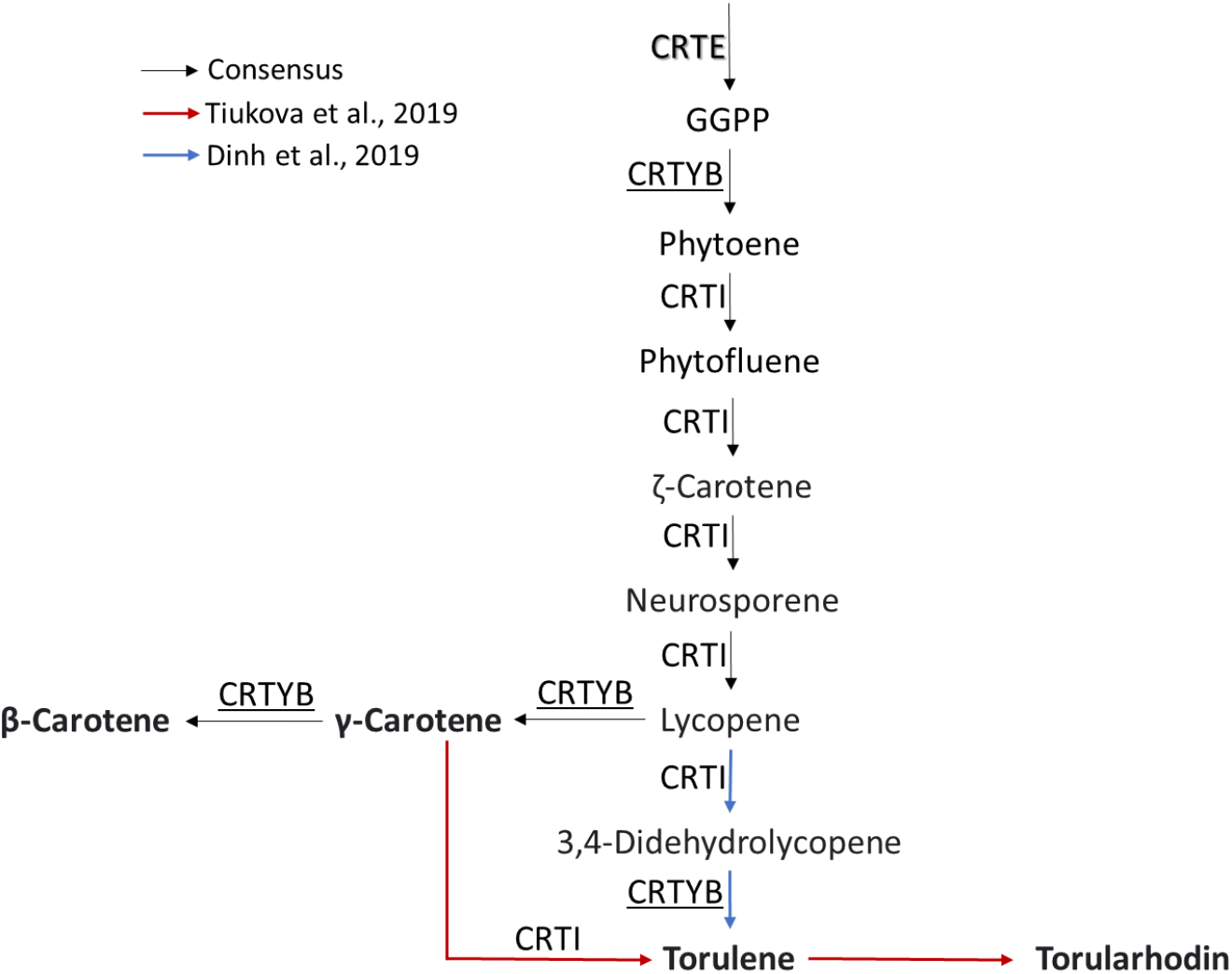
Carotenoid biosynthesis in *R. toruloides* according to different genome-scale models (GEM) (Dinh et al., 2019; Kim et al., 2021; Tiukova et al., 2019). Geranylgeranyl diphosphate synthase (crtE) catalyzes the formation of precursor of carotenoids. Phytoene dehydrogenase (crtI) and phytoene synthase/lycopene cyclase (crtYB) are both involved in the production of, respectively, acyclic and cyclic carotenoids.

When compared to the parental strain (*R. toruloides* CCT7815), only SBY92 showed differences in biomass production (Supplementary Figure S1). The xylose consumption profile (Supplementary Figure S1) and the maximum specific growth rates (µ_max_) were identical for all strains (*R. toruloides* CCT7815, and SBY93: 0.12 ± 0.00 1/h and SBY94: 0.12 ± 0.01 1/h), except for SBY92 which was statistically significant slightly lower (µ_max_: 0.11 ± 0.00 1/h). According to the quantification done with the HPLC method (same used for previous publications on *R. toruloides* CCT7815), the parental strain produced 9.9 ± 0.2 mg/l, while SBY92-94 produced 11.9 ± 1.3, 14.0 ± 0.3, 13.8 ± 0.5 mg/l of total carotenoids, respectively (Figure 4A). Concentrations measurements using the spectrophotometer were on average 41% higher than the ones using the HPLC method, probably due to the presence of other intermediate carotenoids that were not detected with the latter.

**Figure 4.**
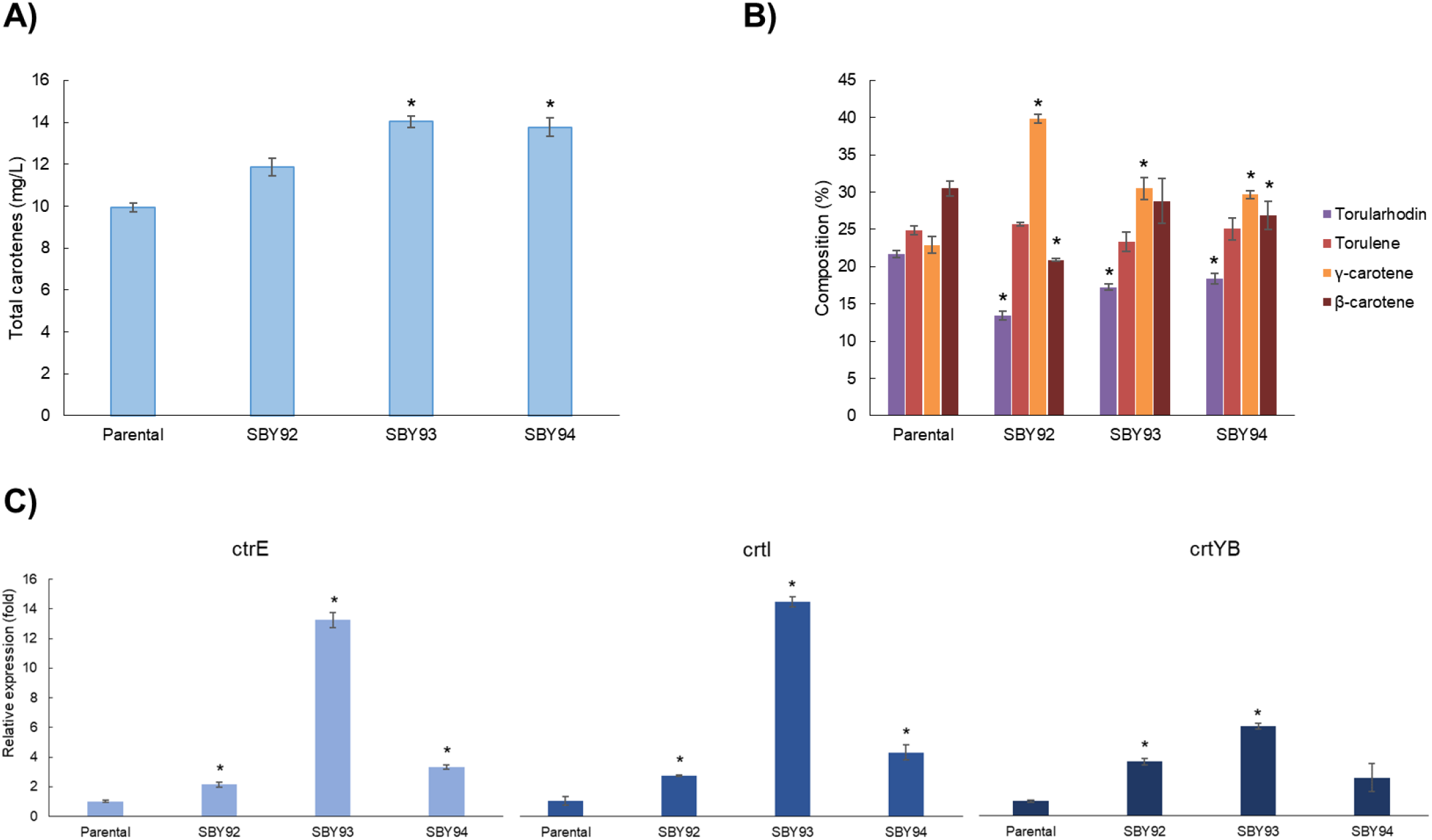
**A)** final carotenoid titer and **B)** composition (HPLC method). **C)** Relative expression of the genes crtE, crtI, and crtYB in the strains SBY92-94 in comparison to parental strain during the exponential phase. Asterisk shows the significative statistical difference (Student’
ss t-test; p-value ≤ 0.05) when compared to the parental strain (*R. toruloides* CCT 7815).

SBY92 was the only modified strain that showed a statistically significant improvement in carotenoid yield on biomass (Y_X/CAR_: 2.01 ± 0.06 mg carotenoid/g dry cell mass) when compared to the parental strain (Y_X/CAR_: 1.23 ± 0.04 mg carotenoid/g dry cell mass), probably due to the lower achieved biomass (Supplementary Figure S1). Regarding carotenoid composition (Figure 4B), all modified strains showed lower torularhodin and increased γ-carotene percentages in the total carotenoid composition than the parental strain. Moreover, the SBY94 showed also, an increased β-carotene percentage. SBY92-94 also showed higher expression of crtE, crtI, and crtYB, validating the overexpression of the pathway genes (Figure 4C).

A Spearman correlation (ρ) was run to assess the monotonic relation between the expression of crtE, crtI, and crtYB and carotenoid titers. The correlations were very strong for crtE and crtI (ρ; crtE: 0.88; crtI: 0.93; p-value; crtE: 0.004; crtI: 0.001; n: 8) and strong for crtB (ρ: 0.76; p-value: 0.028; n: 8). The non-linearity of these correlations could be related to bottlenecks in the supply of geranylgeranyl diphosphate (GGPP) precursors (Figure 3). In this case, an increase in the expression of the carotenogenic genes would be beneficial only until the available crtE and crtI are saturated with their substrates. After this point, there is an excess enzyme, but the amount of active enzyme remains the same, as does the maximum rate of reaction. Considering that SBY94 relative expressions of crtE, crtI, and crtYB are, respectively, 4.0, 3.4, and 2.4 lower than SBY93 (Figure 4C), yet the carotenoid titer produced by these strains is statistically the same (Figure 4A), we suggest that expression levels higher than those presented by SBY93 would no longer be beneficial without the overexpression of genes upstream of crtE. Genes from the mevalonate pathway, such as HMG-CoA reductase, are recommended targets for overexpression to increase carotenoid production (Tiukova, 2019) and have already shown positive results for this strategy for other carotenogenic yeats (Wang and Keasling, 2002).

The relative expression of crtE, crtI, and crtYB genes may also be affecting the distribution between carotenoids derived from lycopene; γ-carotene and β-carotene, and 3,4-dihehydrolycopene; torulene and torularhodin (Figure 3). This classification assumes that γ-carotene is not converted to torulene by crtI (Dinh et al., 2019; Hausmann and Sandmann, 2000; Schaub et al., 2012). SBY92 and SBY93 have accumulated proportionally more γ-carotene and β-carotene than parental strain and SBY94 (Table 2). Lycopene is both substrate and inhibitor of lycopene cyclase, present in crtYB (Ma et al., 2022). The inhibition increases and β-carotene selectivity decreases proportionally to the lycopene rate of formation (Ma et al., 2022), which is mediated by crtI (Figure 3). Therefore, selectivity towards γ-carotene/β-carotene is proportional to the active crtYB/crtI ratio. Those were calculated for *R. toruloides* CCT7815 and SBY92-94 using their respective relative expression values (Table 2; Figure 4C). For this calculation, we considered it reasonable to use SBY94’s relative crtI expression also for SBY93’s ratio, due to the reasons discussed in the previous paragraph. The calculation results were consistent with what was observed experimentally: SBY92 and SBY93 have higher crtYB/crtI ratios and γ-carotene/β-carotene selectivity.

**Table 2.** Carotenoid titer and genes ratios for *R. toruloides* CCT7815 and SBY92-94.

| Strain | γ-car + β-car/trl+trlhd | crtYB/crtI |
| --- | --- | --- |
| parental | $1.15 \pm 0.014^*$ | $1.03 \pm 0.34$ |
| SBY92 | $1.55 \pm 0.025^\dagger$ | $1.34 \pm 0.08$ |
| SBY93 | $1.46 \pm 0.101^\dagger$ | $1.43 \pm 0.03^\ddagger$ |
| SBY94 | $1.30 \pm 0.100^*$ | $0.58 \pm 0.21^\dagger$ |

$\gamma$ -car: γ-carotene; $\beta$ -car: β-carotene; trl: torulene; trlhd: torularhodin
\*: statistically the same
$^\dagger$ : statistically the same
$^\ddagger$ : calculated with crtI ratio of SBY94

The concentrations for carotenoids produced by *R. toruloides* grown on glucose range from 14.0 to 33.4 mg/l (Dias et al., 2015). *R. toruloides* CCT7815 has been reported previously to produce 1.1 and 1.72 mg/l of total carotenoids in xylose-rich hydrolysates from sugarcane bagasse and birch, respectively (Bonturi et al., 2017b; Oliveira et al., 2021). In pure xylose cultivation in bioreactors, this strain was able to produce 20.7 and 36.2 mg/l without and under light irradiation (Pinheiro et al., 2020). For *R. glutinis* cultivated in brewery effluent, the literature reports 1.2 mg/l total carotenoids (Schneider et al., 2013). By optimizing the fermentation conditions, changing C/N ratios, and doing several rounds of metabolic engineering, *Y. lipolytica* was able to produce 33 mg/g dry cell mass of carotenoids (Gao et al., 2017). Using optimum promoter-gene pairs for heterologous carotenoid production pathway, Larroude et al. (2017) were able to engineer a strain of *Y. lipolytica* with a maximum yield of 90 mg/g dry cell mass of β-carotene. Literature uses either HPLC or spectrophotometric methods to quantify carotenoids but results are divergent. Liu et al., (2021) showed even when using the same methodology different carotenoids as standards led up to 40% variation in the results. Therefore, it is suggested the development of a standard methodology to allow a fair comparison between different strains and studies from different groups. The titers of total carotenoids achieved in this work are superior or within the expected margin of concentrations attained in non-optimized strains and cultivation conditions. However, to transform *R. toruloides* into a competitive biotechnological producer of carotenoids, it is necessary to further fine-tune the expression of the carotenoids synthesis pathway, elimination of competing pathways, and as well as optimize the cultivation conditions.

## 4 Conclusions

In this work, the existing collection of YTK for different yeasts was adapted to *R. toruloides*, requiring optimization of almost all steps of the assembly, which is, to our knowledge, the first dedicated GGA for a basidiomycete. The RtGGA platform was further validated by the assembly of a construct consisting of three genes, a selection marker, and insertional units. The three genes, coding for essential proteins in the carotenoids biosynthesis pathway were overexpressed and led to increased carotenoid production (up to 41%). Increased gene expression was confirmed by RT-PCR. The establishment of a golden gate assembly platform for *R. toruloides* fills a gap of advanced genome engineering tools for this non-conventional yeast that has engineering potential for industrial biotechnology applications. Optimization of cultivation conditions to increase carotenoid yield, as well as using the now established GGA platform to help with overexpressing key proteins in the lipid production pathway can make *R. toruloides* an industrial yeast capable of producing fine chemicals and biofuels, helping in the transition towards a sustainable economy.

## Authors Contribution

NB, MJP, RLA, P-JL designed the experiments. NB, MJP, ER, PMO, GL, and TE performed the experiments. AB and MR performed the assembly and annotation of the genome. NB, MJP, PMO, GL, EAM, RLA, P-JL analyzed the data. NB, ER, JSDB, and MJP wrote the manuscript. All authors revised the manuscript.

## Acknowledgments

We thank the Institute of Genomics at the University of Tartu for the genome sequencing, Professor Mart Loog’s lab for kindly providing the parts from iGEM Collection, Laura Kibena and Andréia Axelrud for the help with HPLC analysis, Astrid Sallumäe for experimental help, and Solis BioDyne for providing tailored Taq polymerases.

---

**Supplementary Figure S1.**
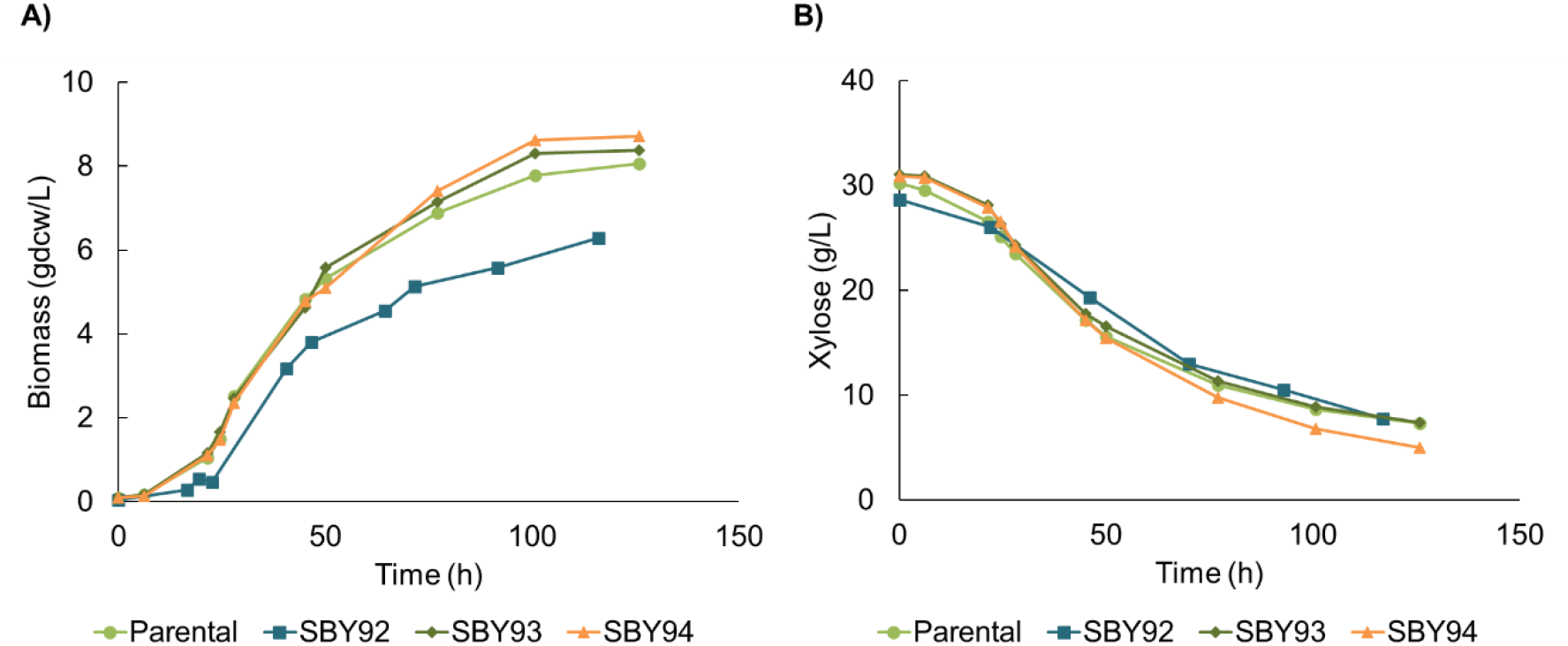
**a)** Growth curve and **b)** xylose consumption profile of strains *R. toruloides* CCT7815 (parental) and SBY92-94. Errors were calculated in terms of standard deviation (n=3) and were ≤15%.

**Supplementary Table 1.**
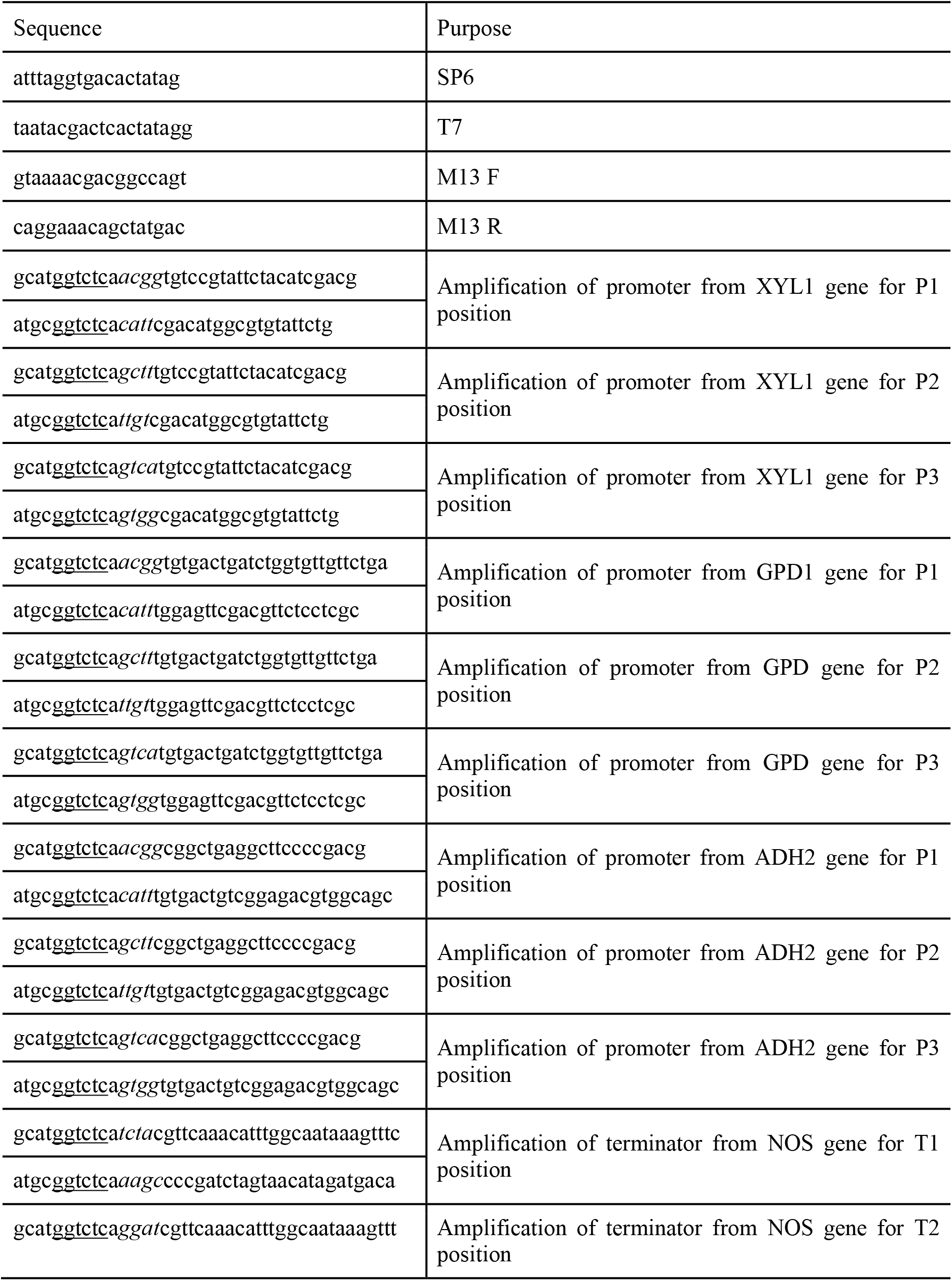
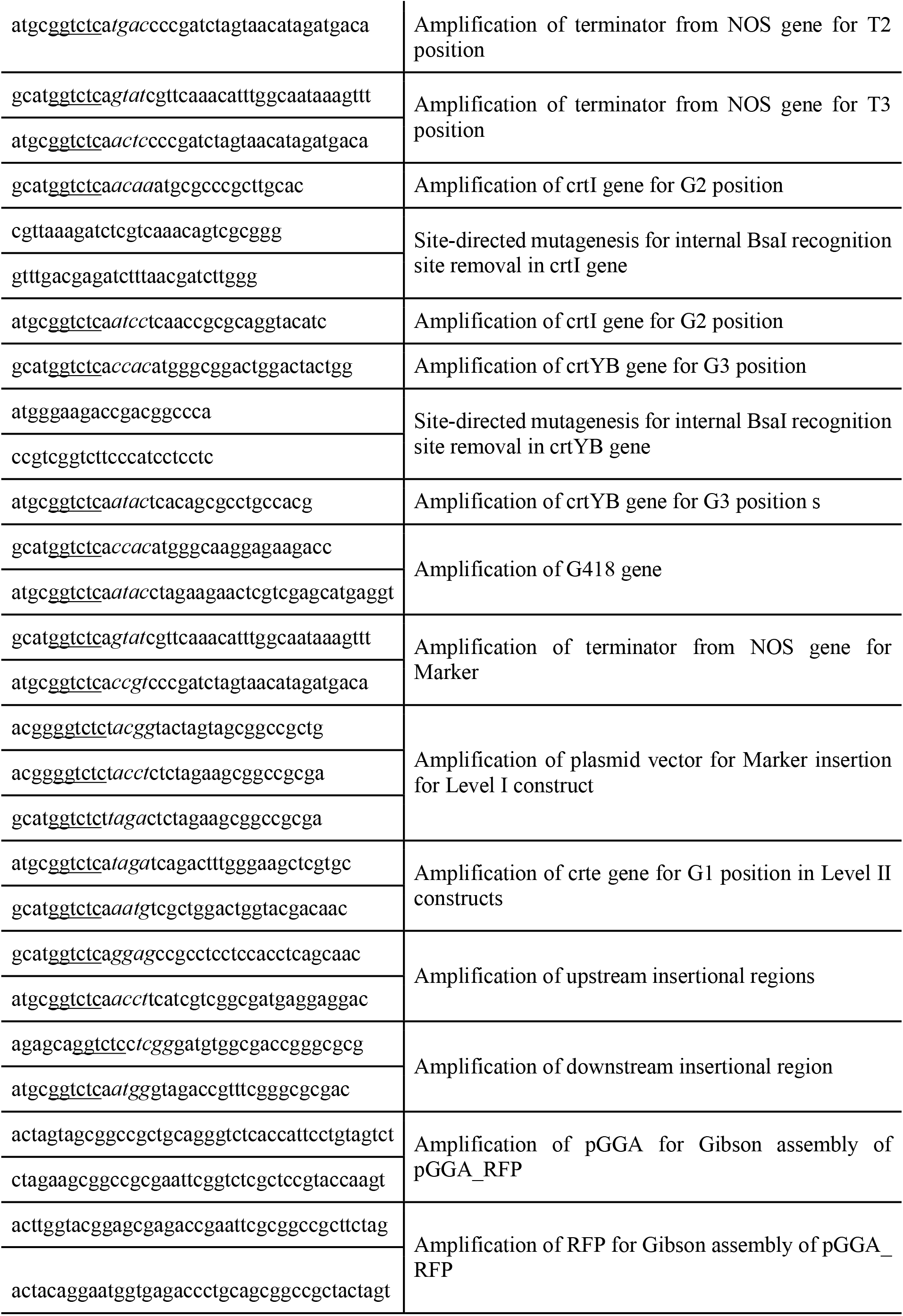
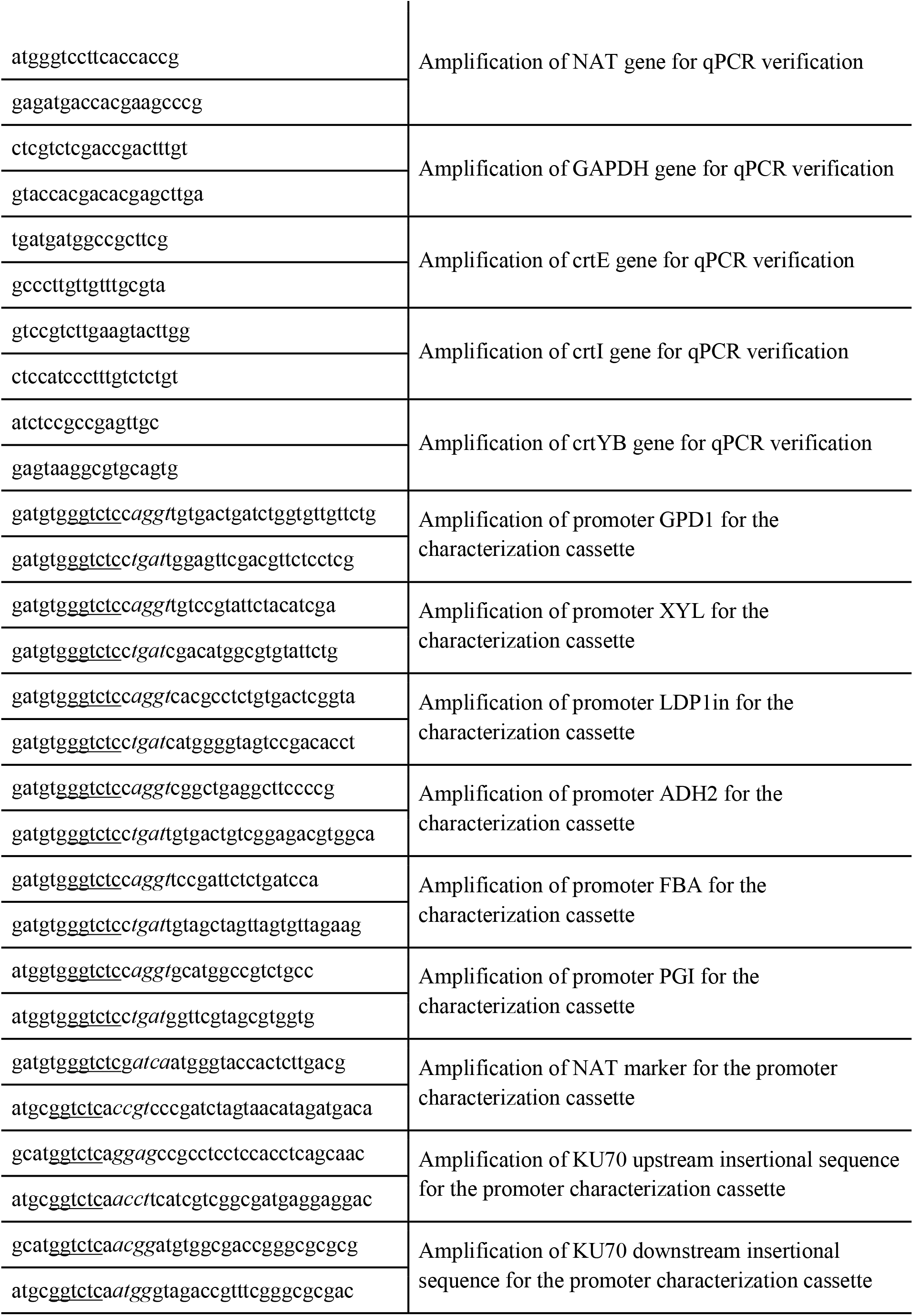
A list of oligonucleotides used for Golden Gate assembly. In the ‘Sequence’ column, the lowercase base pairs are for enzyme anchoring, BsaI recognition site is <u>underlined</u>, and the 4-base pair overhangs are in *italic*.

